# Low-Energy Polishing Facilitates Breaking the Resolution Barrier of In Situ Cryo-EM

**DOI:** 10.64898/2026.06.09.731061

**Authors:** Chunling Wu, Qi Yang, Xiaodong Su, Mei Li, Xinzheng Zhang

## Abstract

In situ structural analysis allows direct visualization of protein structures in their native cellular environments, but near-atomic resolution in cellular lamellae has been significantly limited to exceptionally large complexes such as ribosomes. A key factor underlying this limitation is the degradation of data quality of thin lamellae caused by substantial subsurface damage from cryo-focused ion beam (cryo-FIB). Here, we developed a cryogenic low-energy polishing in FIB approach, which reliably produces thin lamellae with low damage across different cell types. This advance, combined with in situ single particles analysis, has pushed down the molecular weight lower limit for in situ reconstruction at near-atomic resolution to ∼400 kDa. We demonstrate this by resolving photosynthetic complexes (3.4 Å and 3.3 Å), metabolic enzymes (3.3 Å), chloroplast ribosome (4.0 Å) and respiratory chain complexes (3.7 Å) from *Chlamydomonas reinhardtii*. Furthermore, the efficient workflow enables rapid structural feedback upon changes in cellular states, offering a practical way to perform multi-condition in situ structural analysis.

## Introduction

Cryo-electron microscopy (cryo-EM) is one of the key techniques advancing structural biology. For in vitro studies, single-particle analysis (SPA) has matured into a powerful technology. The “resolution revolution” ^1,2^ propels structural determination from medium to near-atomic resolution (sub-4 Å), enabling detailed atomic modeling and driving rapid advances in the field. In recent years, increasing efforts have been directed toward the development of in situ structural analysis methods. Cryo-electron tomography (cryo-ET) currently serves as the primary technique for in situ structural studies ^3,4^. With the development of subtomogram averaging (STA) software ^5–9^, the resolution of in-situ structural determination has been significantly improved ^10–14^. However, for cryo-lamellae, structural determination of most proteins remains constrained to 5–8 Å mainly limited by time-consuming tilt-series acquisition and inaccuracy alignment of tomogram ^15,16^. Currently, only ribosomes routinely achieve near-atomic resolution, largely due to their high cellular abundance and large molecular weight ^8,17–19^. The structural analysis of membrane proteins is even more difficult due to their extensive transmembrane domains which are embedded in the native membrane ^20,21^. Thus, the transmembrane domains hardly contribute to particle picking and the initial alignment in STA, which limited the resolution of membrane proteins with small soluble domains. For example, Photosystem II (PSII) resolved by cryo-ET typically reaches only ∼20 Å resolution^14,20^, while Photosystem I (PSI) has not yet to be visualized in tomogram due to its even smaller hydrophilic domains.

Recently, a new method of in situ single-image analysis has emerged as a complementary approach, such as in situ single-particle analysis (isSPA) ^22,23^ and two-dimensional template matching (2DTM) ^24,25^. Unlike cryo-ET, which relies on tilt-series acquisition, these methods directly achieve high-resolution reconstructions directly from tilt-free micrographs by localizing target proteins using corresponding projections derived from high-resolution 3D templates, significantly improving the throughput of in situ structural analysis. In micrographs of cellular lamellae, densities from surrounding biomacromolecules overlap with those of the target protein and are treated by isSPA as background noise. To improve target localization, an isSPA weighting function is incorporated into cross-correlation (CC) calculation to suppress the effects of both background and shot noises. In addition, isSPA incorporates a special refinement and reconstruction workflow tailored to the background-noise characteristics of cellular lamellae and employs several strategies to minimize false positive detections during reconstruction, thereby improving the achievable resolution. However, the detection efficiency is highly sensitive to the image quality of the lamellae ^23,26,27^ and drops sharply for small proteins, which has limited its broader application. Using thin lamellae to reduce the background noise is a possible way to enhance the detection efficiency.

Cellular or tissue specimens are too thick and must be thinned to nanometer-scale thickness for transmission electron microscopy (TEM) imaging. Cryo-focused ion beam (cryo-FIB) milling is widely used because it offers a high degree of automation and minimal sectioning artifacts ^28–31^. However, studies have shown that conventional 30 keV gallium (Ga^+^) FIB milling causes damages up to 50-60 nm from each biological lamella surface ^26,27^, a critical factor previously underestimated ^29,32^. Consequently, a lamella thickness of 150–180 nm is recommended ^23,33,34^. As a result, the detection efficiency of isSPA no longer improves when lamellae are thinner than 160 nm, and the minimum molecular weight that can be resolved is limited to approximately 1.1 MDa ^23^. Further thinning may cause damaged proteins to dominate in the lamellae. Therefore, minimizing surface damage is essential to achieve high-quality thin lamellae. Reducing the ion-beam accelerating voltage is an effective way to mitigate milling-induced damage in materials science ^35–38^. Marko et al. provided an early description of the use of low-energy ion-beam milling in frozen-hydrated specimen thinning, demonstrating that different milling voltages did not induce detectable heat-induced devitrification ^28^. Subsequently, our previous study showed that lowering the milling voltage to 8 keV in cryo-FIB milling substantially reduces damage depth to 30 nm. However, unlike material samples, biological specimens must be thinned into flat lamellae with uniform thickness at cryogenic temperatures, which imposes persistent limitations on the practical application of low-energy FIB milling in biological specimen preparation. Reducing the acceleration voltage drastically diminishes ion penetration depth and worsens beam convergence ^39,40^, which not only prolongs milling times and reduce processing efficiency, but also makes common lamella-preparation artifacts, such as wedge-shaped thickness variation and curtaining ^30,41,42^, more difficult to suppress. Consequently, obtaining intact and uniform thin lamellae becomes more challenging ^26^, thereby limiting the potential advantages for isSPA localization.

Here, we developed a cryo low-energy polishing in cryo-FIB (cryo-LEP-FIB) approach that reliably produces high-quality thin lamellae across diverse cell types. Our analysis shows that high-quality thin lamella significantly lowers the molecular weight limit of in situ structural determination by isSPA at near-atomic resolution to ∼400 kD. Using this approach, we obtained near-atomic resolution in situ structures of several representative cellular complexes from *Chlamydomonas reinhardtii* (*C. reinhardtii*), including membrane proteins PSII, PSI and mitochondrial respiratory chain complexes, as well as the soluble protein Rubisco and the chloroplast ribosome. In addition, we capture the transient association of photoprotective protein, illustrating its capacity for high-efficient multi-condition structural studying in situ.

## Results

### Low-energy polishing preparing high-quality cellular lamellae

In cryo-LEP-FIB, cryo-samples are first rough-milled at 30 keV Ga^+^ to approximately 300 nm to maintain milling efficiency, followed by a switch to 8 keV for the low-energy polishing step to minimize ion-beam-induced damage (Figure 1A and B). Under conventional 30 keV cryo-FIB conditions, protective organometallic Pt layer deposited by gas-injection-system (GIS)^41,42^ and over/under-tilt milling ^30,43,44^ have been shown to effectively suppress curtaining and reduce wedge effect. However, when the polishing voltage is lowered, direct transfer of these parameters leads to severe breakage at the leading edge (Figure S1A), pronounced front-to-back thinning gradient (Figure S1B), and significantly aggravated curtaining (Figure S1C). To enable robust and reproducible lamella thinning at 8 keV, we optimized three key parameters: doubling the Pt layer thickness, applying a larger symmetric over-tilt of 0.6–0.8° during 8 keV polishing, and increasing the Z-axis milling depth by 5- to 8-fold (details described in STAR Methods, Figures 1C-E and S1). Under low-energy conditions, the doubled Pt layer both protects the leading edge from overmilling and suppresses curtaining caused by Pt nonuniformity, while preserving milling efficiency (Figure S1A). The larger over/under-tilt improves thinning at the trailing region while maximizing the usable lamella area. (Figure S1B). Moreover, the shallower penetration depth at low voltage amplifies local milling-rate variations caused by specimen heterogeneity ^45,46^, making a greater Z-axis milling depth necessary to extend the milling path and progressively smooth curtaining stripes (Figure S1C). Together, these optimizations ensure uniform thickness and preserve lamella integrity, thereby significantly enhancing the success rate of lamella preparation using the cryo-LEP-FIB method.

**Figure 1.**
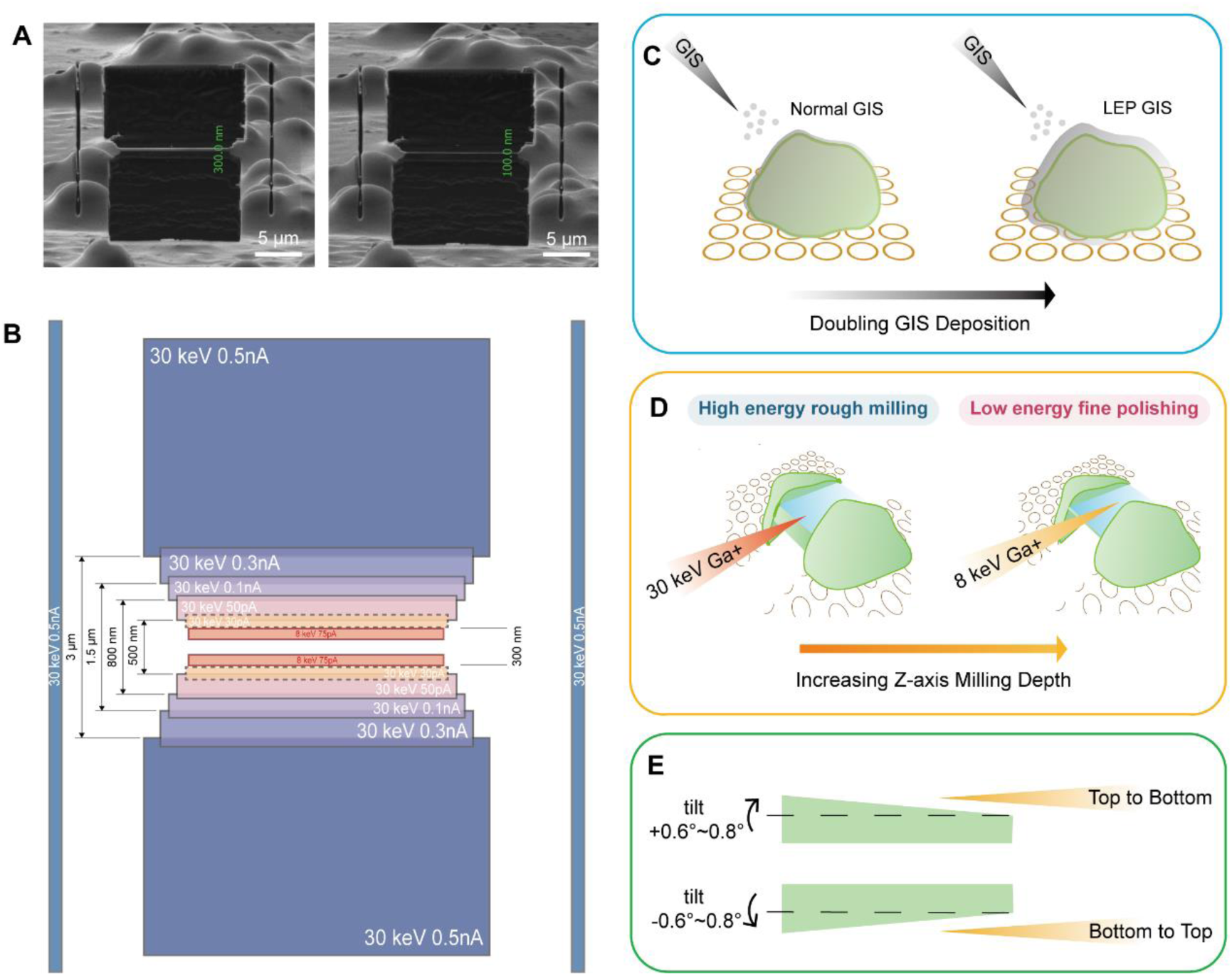
Schematic of the cryo-LEP-FIB milling process. (**A**) Ion-beam image of a representative lamella thinned to ∼300 nm using 30 keV Ga⁺ and further polished to ∼100 nm at 8 keV Ga⁺. (**B**) Voltage and current applied stepwise during cell milling, along with the corresponding lamella thickness, shown for reference. Colored boxes indicate the relative pattern size at each thinning step. (**C**) Prior to milling, the GIS protective layer was enhanced by doubling the deposition time compared with standard 30 keV conditions. (**D**) Compared with 30 keV milling (red beam), the Z-axis milling depth was significantly increased at 8 keV milling (orange beam). (**E**) Cartoon illustrating the effect of increasing over-tilt angle. Lamellae are shown in green, low-voltage ion beam in orange, and dashed lines indicate milling direction.

We successfully validated the cryo-LEP-FIB approach on *C. reinhardtii* cell using a cryo-FIB scanning electron microscopy (SEM) Aquilos 2 microscope (Fig. 2A). ∼20 high quality lamellae can be prepared per day with a success rate exceeding 85% using 30 keV auto-rough milling and 8 keV manual fine polishing (Figures 2B, C and S3A), consistently yielding high-quality cryo-lamellae with a thickness ranging from 40 to 280 nm, ∼50% thinner than 160 nm (Figure 1E). The resulting thin and uniform lamellae allow clear visualization of major cellular organelles in *C. reinhardtii* at low magnification (Figure 2D). In addition, low-energy milling produces thinner lamellae with larger undamaged regions and improved quality, effectively enhancing the signal-to-noise ratio (SNR) of the cryo-EM images ^33,47^. Using the 750-kDa body of the small subunit (SSU) from *C. reinhardtii* as a template, we successfully localized 80S ribosomes (Figures 2F and S2) and achieved a resolution of 3.7 Å. To demonstrate generality, we also prepared lamellae of *Saccharomyces cerevisiae* (*S. cerevisiae*) and *Escherichia coli* (*E. coli*) by cryo-LEP-FIB (Figure 2A) with a thickness of 43%, 65% thinner than 160 nm (Figure S3B and C), respectively. Using the bodies of SSUs of 750 kDa (*S. cerevisiae*) and 530 kDa (*E. coli*) as templates, we resolved the intact 80S (Figure S4) and the 70S ribosome (Figure S5) at 3.4 Å and 3.5 Å resolution, respectively. These results indicated that that low-energy polishing can effectively improve the detection efficiency of isSPA and expand its applicability to substantially smaller targets well below the previously reported 1.1 MDa limit.

**Figure 2.**
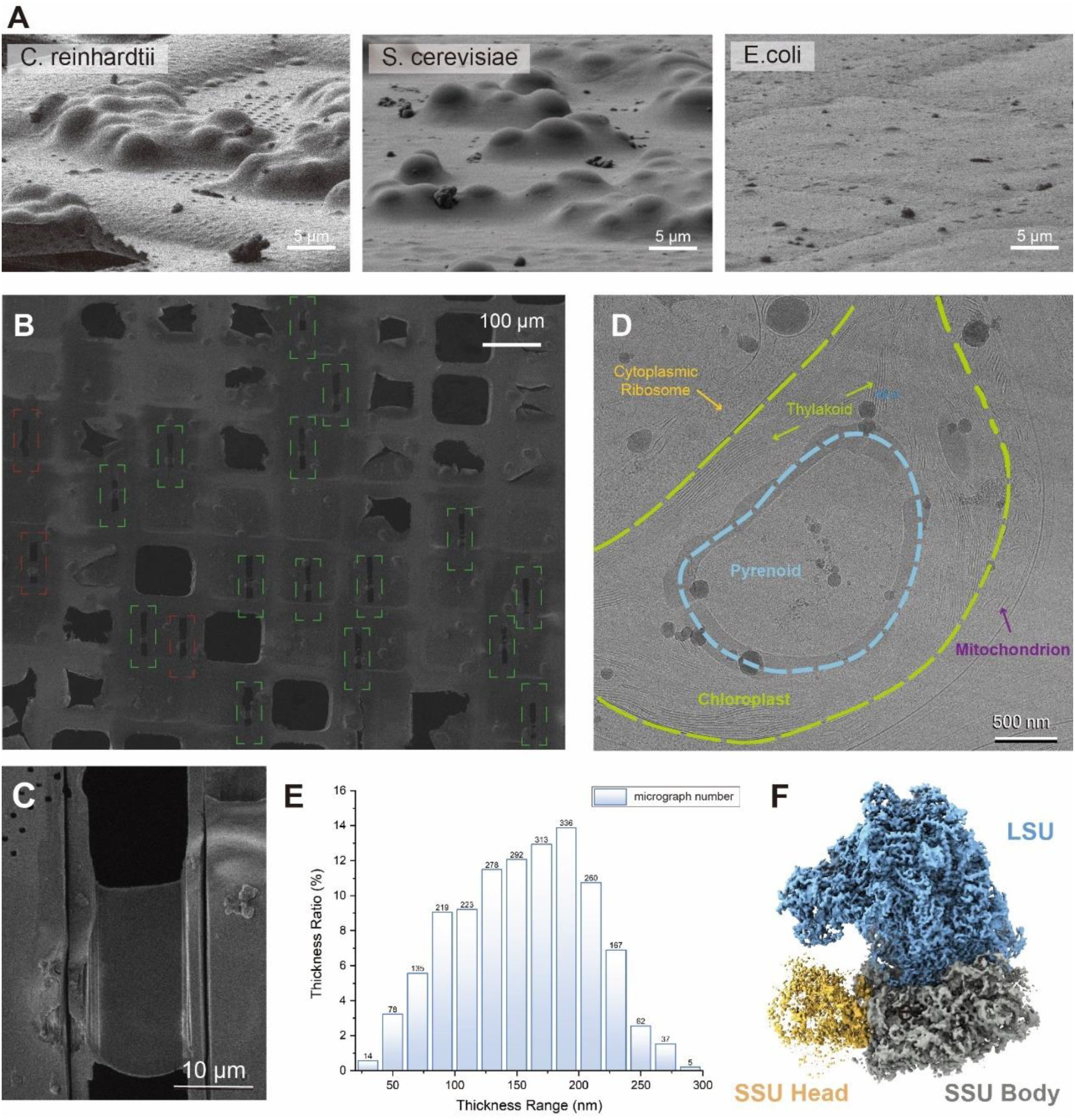
Application and lamellae quality of Cryo-LEP-FIB. (**A**) Ion beam images of three representative cell types prior to thinning. (**B**) SEM overview of *C. reinhardtii* lamellae prepared by cryo-LEP-FIB; intact lamellae are outlined with green dashed boxes, whereas damaged lamellae are indicated with red dashed boxes. (**C**) Enlarged SEM view of a representative *C. reinhardtii* lamella. (**D**) TEM image showing a local view of *C. reinhardtii* cell (scale bar, 500 nm). The chloroplast region is highlighted by a green dashed outline, the pyrenoid is outlined with a blue dashed contour, and representative organelles are indicated by colors. (**E**) Thickness distribution of cell lamellae prepared by Cryo-LEP-FIB. (**F**) Cryo-EM density of the cytoplasmic ribosome detected using the SSU body (grey) as the template. Blue, LSU; yellow, SSU head.

### Thin lamellae extending the molecular weight Limit

Previous studies have demonstrated that reducing the milling voltage to 8 keV can decrease the damage depth to 30 nm ^26^. Based on this, the 60–80 nm thickness range is considered theoretically optimal, accounting for approximately 5% of our total dataset. To define the lower molecular-weight limit accessible for in situ structural determination via cryo-LEP-FIB, we analyzed the precision-recall (P-R) curves for complexes of different sizes in 60–80 nm lamellae using isSPA (Figure 3A and B, see STAR Methods). The P-R curve comprehensively reflects particle detection efficiency; at a given precision threshold, the improvement in recall reflects increased data quality^22,23^. We regard 20% precision as the critical cutoff value for structural determination, as high-resolution reconstruction is easily achieved beyond this threshold through further processing. Complexes ranging from 400 kDa to 1 MDa were generated by clipping from the large subunit (LSU) of the cytoplasmic ribosome of *C. reinhardtii* for simulations of different molecular weights (Figure 3A). At 1 MDa, the recall rate reached 91.5%, indicating near-complete particle detection, and remained high at 73.0% for 600 kDa complexes (Figure 3B). Notably, even at 400 kDa, a recall of 37.1% was achieved. Further validation was performed in both *S. cerevisiae* and *E. coli* samples. At the same precision threshold, recall rates of 27.0% and 24.4% were achieved respectively at 400kDa, demonstrating comparable detection capability for small proteins in cryo-LEP-FIB lamellae (Figure S6).

**Figure 3.**
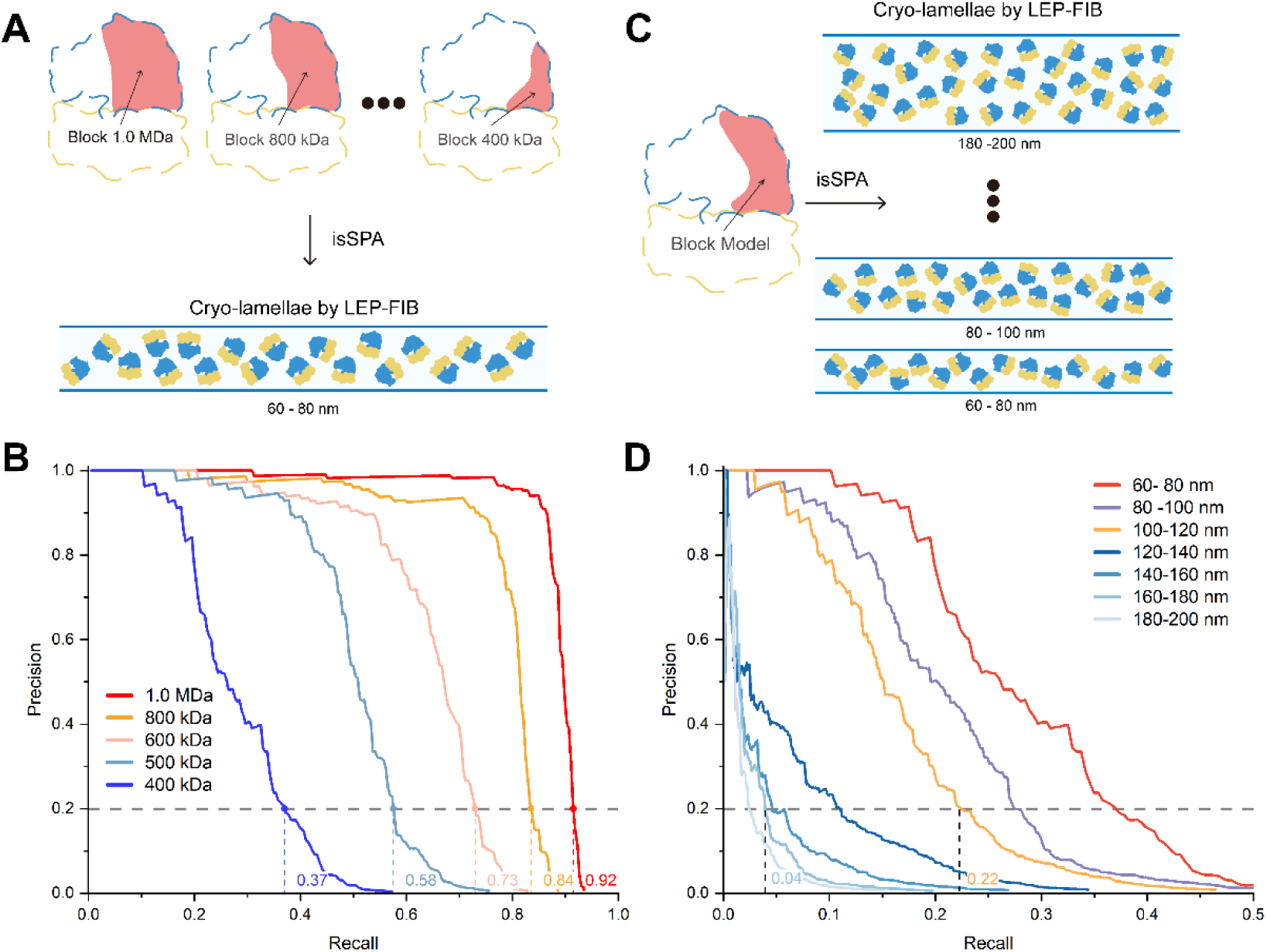
P-R curves analysis of the *C. reinhardtii* cytoplasmic ribosome. (**A**) Schematic illustration of the P-R curve analysis shown in B, in which LSU blocks of different molecular weights were used as templates to detect particles within lamellae of defined thickness. (**B**) Corresponding P-R curves for LSU blocks of varying molecular weights in lamellae 60-80 nm thick, with different molecular weights indicated by colors. Each curve represents data averaged from 10 micrographs. (**C**) Schematic illustration of the P-R curve analysis shown in D, in which a target of defined molecular weight was detected within lamellae. (**D**) Corresponding P-R curves for a 400 kDa target, with lamellae grouped in 20 nm thickness intervals from 60 to 200 nm, indicated by colors. Each curve represents data averaged from 10 micrographs.

To determine the mechanistic basis of this improved detectability, we calculated the detection efficiency at different depth layers from the lamella surface based on raw data previously reported in ^26^, which were reprocessed for the present analysis (Figure S7, see STAR Methods). In lamellae milled at 30 keV, detection efficiency decreased monotonically toward the surface within the first 60 nm, with recall falling below 2% in the outermost 20 nm, consistent with severe damage reported previously ^26,27^. By contrast, lamellae thinned at 8 keV exhibited a plateau in detection efficiency beyond 30 nm depth. At depths of 20 nm and 30 nm, detection efficiency was increased by 8.2-and 3.7-fold, respectively, compared with 30 keV milling. These results indicate that reduced ion beam-induced damage plays a central role in extending in situ structural analysis toward low-molecular weight complexes.

We next assessed how total lamella thickness constrains the detectability of small protein complexes by performing P-R curve analysis using the 400 kDa block across different thickness groups (Figure 3C and D, see STAR Methods). Overall, detection efficiency improves as lamella thickness decreased from 220 nm to 60 nm (Figure 3D). Importantly, the recall rate remained above the 20% within the 100-120 nm thickness range, indicating that lamellae thickness less than 120 nm (28%) are sufficient for reliable localization and structural analysis of ∼400 kDa targets. Further increases in thickness led to a sharp decline in recall, substantially reducing analysis efficiency. Together, these results demonstrate that low-damage, thin lamella is critical for extending in situ structural determination to small proteins complexes by isSPA.

### High-efficiency and high-resolution in-situ structural determination

As a unicellular model organism, *C. reinhardtii* provides a suitable system for testing in situ structural analysis ^14^. The chloroplast of *C. reinhardtii* harbors a highly diverse proteome with molecular weights spanning an order of magnitude, including transmembrane proteins (e.g., PSII, PSI, and respiratory chain complexes), soluble stromal proteins (e.g., Rubisco), relatively low-abundance proteins (e.g., chloroplast ribosome). We performed 8 hours of data acquisition on 12 lamellae prepared via cryo-LEP-FIB, performed isSPA reconstructions, and determined near atomic resolution structures of representative proteins across a broad molecular weight range within their native cellular environment.

As key components of the cellular proteome, membrane proteins play vital roles in sustaining life. For example, PSII is the sole membrane protein complex that catalyzes light-driven water splitting to generate oxygen and high-energy electrons, underpins photosynthetic energy metabolism ^48^. Despite its importance, its in-situ structural analysis remains a persistent challenge in tomography. In eukaryotic photosynthetic organisms, PSII on the thylakoid membrane contains a dimeric core (C_2_, ∼800 kDa) as its structural foundation. Through association with a peripheral antenna system composed of a variable number of light-harvesting complex II (LHCII) subunits, it ultimately assembles into diverse types of PSII supercomplexes ^49–51^. We performed structural determination by using the C_2_ (see STAR Methods) as the template, and successfully resolved the structure of the C_2_S_2_M_2_L_2_ (strongly, moderately and loosely bound LHCII trimers) type PSII supercomplex at a resolution of 3.4 Å (Figures 4 and S8). Compared to purified structure (EMD-9956) ^52^, where the PsbP and PsbQ subunits are prone to dissociation, our structure (Figure 4A and B) clearly reveals that both subunits are stably anchored to the luminal side of the PSII core. Additionally, PsbY and PsbR, which were not found in the structure of purified C_2_S_2_M_2_L_2_ type PSII, are observed to bind to the periphery of the supercomplex in the thylakoid membrane. Notably, we identified a previously uncharacterized helical density near CP47 on the luminal side of the core complex (Figure S9), which we refer to as the unknown luminal protein (ULP). In addition, P-R curve analysis of PSII revealed that the detection efficiency of the membrane protein is comparable to that of water-soluble proteins (Figure S10A and B). To further verify the molecular weight limit of cryo-LEP-FIB, using the monomeric core (400 kDa) as a template, we also successfully determined the C_2_S_2_M_2_L_2_ type PSII supercomplex (Figure S10C-F).

**Figure 4.**
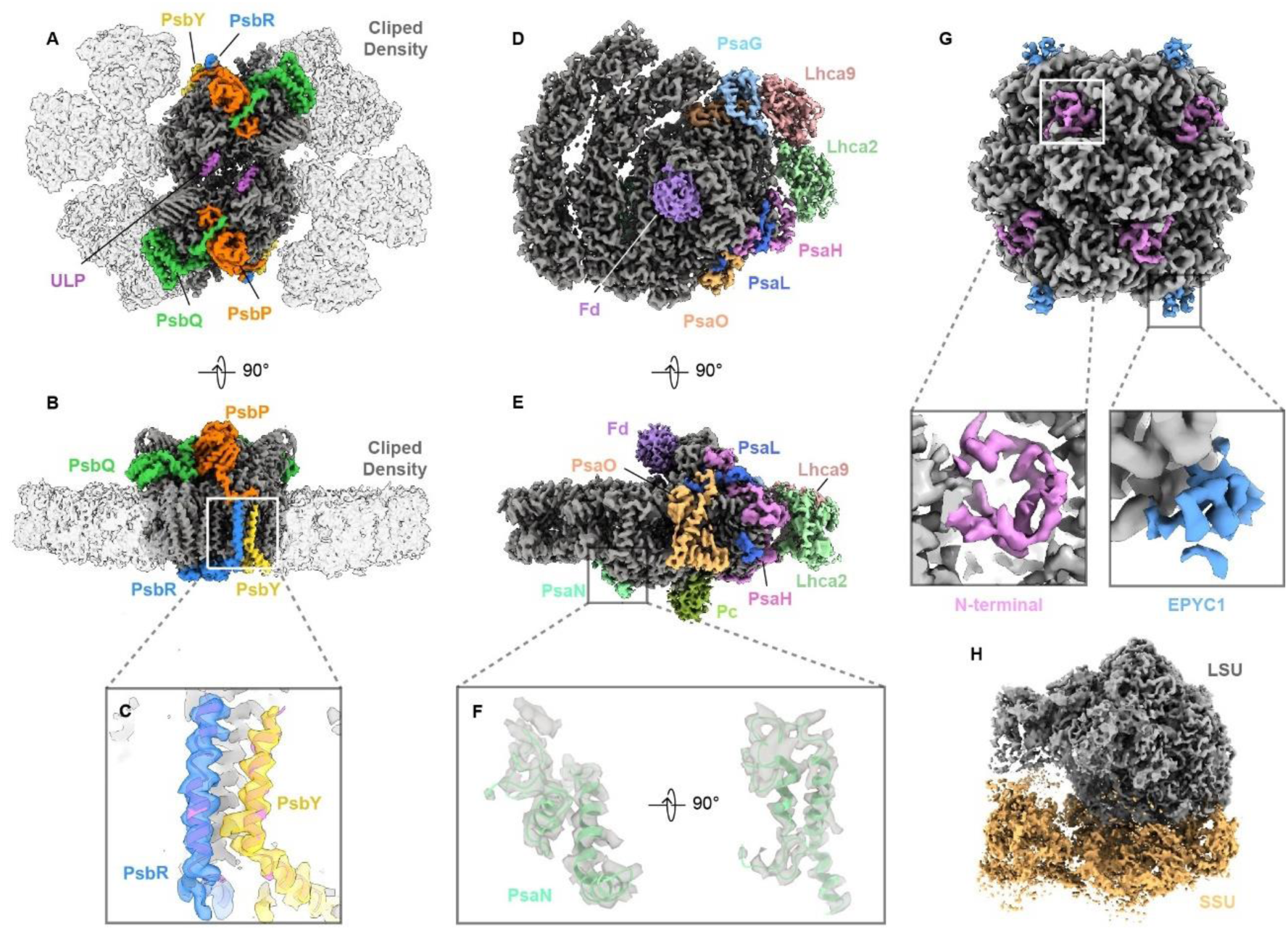
High-resolution in situ structures of representative complexes in *C. reinhardtii*. (**A**) Overall densities of the C_2_S_2_M_2_L_2_ supercomplex shown from the stromal side. The template used for particle detection is colored dark grey, with light-harvesting antennae densities (light grey, transparent) excluded from the SPA structure (EMD-9956). Differences from the purified SPA structure are highlighted in distinct colors. (**B**) C_2_S_2_M_2_L_2_ supercomplex viewed from the membrane side (left) and (**C**) closed-up view of well-resolved densities of PsbR (blue densities) and PsbY (yellow densities). (**D,E**) Cryo-EM density of the PSI supercomplex shown from (**D**) stromal side and (**E**) membrane side. The template is colored dark grey, with differences from the purified structure (EMD-9678) shown in distinct colors. (**F**) The closed-up view of well-resolved densities of PsaN in membrane and stromal sides (left and right), with the model colored in green. (**G**) In situ structure of Rubisco (top). The particle-detection template is shown in dark grey; close-up views (bottom) highlight the N-terminal region of the large subunit (LSU, pink) and EPYC1 (blue). (**H**) In situ structure of chloroplast ribosomes.

PSI captures light energy and drives electron transport toward NADPH synthesis, functioning alongside PSII as an indispensable dual engine of photosynthetic energy conversion ^53^. Using the PSI complex (EMD-9678) ^54^ as a template (775 kDa), we resolved the in situ structure of PSI at a resolution of 3.3 Å (Figure 4, Figures. S11 and S12). The in situ structure fully resolves the native PSI assembly complex, including the intact Lhca2, Lhca9, PsaG, PsaO, PsaL, PsaH, and PsaN subunits, which are often missing in purified complexes. Importantly, plastocyanin (Pc) and ferredoxin (Fd)—electron carriers that often dissociate during purification—remained bound to the PSI core in their native states. This allowed us to visualize the complete electron transport chain within PSI. In addition to the PSI complex, the classification also yielded 25% PSI-LHCII supercomplex with an overall resolution of 3.8 Å, while the LHCII domain reached 4.7 Å (Figures. S11 and S12).

In addition, we resolved the structure of the RuBisCO enzyme within the chloroplast pyrenoid. RuBisCO is the key enzyme in carbon fixation, catalyzing the conversion of atmospheric CO₂ into organic molecules. In *C. reinhardtii*, RuBisCO molecules form a dense, phase-separated matrix within the pyrenoid, which enhances catalytic efficiency by concentrating CO₂ around the enzyme ^55,56^. Despite the high packing density of RuBisCO within the pyrenoid, we achieved a in situ reconstruction at 3.2 Å resolution (Figures 4G and S13). The RuBisCO holoenzyme adopts a conformation consistent with the reported structure ^57^, comprising catalytic LSUs and SSUs that stabilize the assembly. Compared to the SPA-determined structures, the large subunit exhibits a more complete N-terminal domain. Moreover, we identified eight additional densities bound to the small subunits in situ (Figure 4G), whose structure and spatial arrangement match those of Essential Pyrenoid Component 1 (EPYC1) ^58,59^—a key factor in pyrenoid formation—providing further structural evidence for the phase separation mechanism.

Chloroplast ribosomes specifically translate mRNAs encoded by the chloroplast genome, directly synthesizing the core subunits of key photosynthetic components—including the reaction center proteins of PSI and PSII, as well as the large subunit of RuBisCO—which serve as the structural foundation for photosynthetic function. However, the abundance of chloroplast ribosomes is significantly lower than that of cytosolic ribosomes. Using the large subunit of chloroplast ribosome (see STAR Methods) as a template for particle detection, we identified only 11,588 high-confidence particles from 891 micrographs, averaging 13 particles per micrograph, corresponding to just one-twentieth of the abundance typically obtained for cytoplasmic ribosomes ^8,18,19,60^. From this dataset, we reconstructed the LSU region at 4.0 Å resolution (Figures 4H and S14). Nevertheless, the current particle number is insufficient, limiting the ability to resolve high-resolution functional conformations of the chloroplast ribosome during translation.

In addition to chloroplasts, mitochondria play a central hub of energy production in eukaryotic cells, where oxidative phosphorylation is carried out by the respiratory chain embedded in the inner membrane. Respiratory complexes I, III, and IV (CI, CIII and CIV) assemble into functional groups that couple electron transfer to transmembrane proton translocation, generating the proton motive force that drives ATP synthesis and sustains cellular physiology ^11,61^. However, the native organization and functional states of respiratory chain complexes within intact mitochondria remain insufficiently characterized. In contrast to chloroplasts, which occupy most of the cellular volume, mitochondria are comparatively sparse in *C. reinhardtii*. To enable high-resolution visualization of mitochondrial respiratory chain complexes, we performed an additional twofold data acquisition. Using a CICIII₂ assembly (EMD-50202) as the template, we first resolved a CI₂CIII₄CIV₆ respiratory chain supercomplex (Figures 5 and S15). Its overall organization is consistent with previous reports ^11^, and the reconstruction reached an overall resolution of 4.1 Å (Figure S15). The CI₂CIII₄CIV₆ supercomplex exhibits a pseudo-C2 symmetry. In the asymmetric unit, CIII₂ is positioned on the concave side of the CI membrane arm in a classic arrangement, and the two CI₂CIII₂ complexes are head-to-head, forming the core scaffold of the supercomplex. Further local refinement of CI and CIII₂ resulted in resolutions of 3.7 Å and 3.8 Å, respectively (Figures S15 and S16), enabling clear visualization of functional transmembrane side chains, reaction-center conformations, and cofactor positions, including the ND2 (NADH dehydrogenase subunit 2) side chain (Figure 5C) and the ubiquinone-10 (Q10)-binding site (Figure 5D) in CI, as well as the high- and low-potential hemes bH and bL in CIII_2_ (Figure 5E). On both sides of the scaffold, three CIV complexes occupy the lateral membrane platforms (Figure 5A and B), referred to as proximal CIV (CIV_P_), middle CIV (CIV_M_), and distal CIV (CIV_D_). The CIV densities show heterogeneous occupancy, prompting focused classification. CIV_M_ is consistently associated with the CICIII₂ complex, forming a fundamental CI₂CIII₂CIV_M_ unit, for which local refinement of CIV_M_ reached 4.1 Å (Figures. S15 and S16). Focused classification of CIV_P_ revealed an approximately 9:1 ratio between bound and unbound populations. Among CIV_P_-bound particles, further classification on CIV_D_ showed that 48.3% contain CIV_D_, yielding a CI₂CIII₂CIV_PMD_ unit (Figures 5F and S15J), whereas in the remaining 42.7%, the CIV_D_ position is occupied by a phospholipid bilayer, resulting in a CI₂CIII₂CIV_PM_ unit (Figures 5F and S15J). Classification of CIV_P_-unbound particles identified units containing both CIV_M_ and CIV_D_ (CI₂CIII₂CIV_MD_; Figures 5F and S15I), as well as units containing only CIV_M_ (CI₂CIII₂CIV_M_; Figures 5F and S15I). Collectively, we captured four distinct compositional states of the CICIII₂CIV_n_ unit. The variability of CIV likely provides the supercomplex with functional flexibility, enabling dynamic regulation of electron transfer and proton translocation.

**Figure 5.**
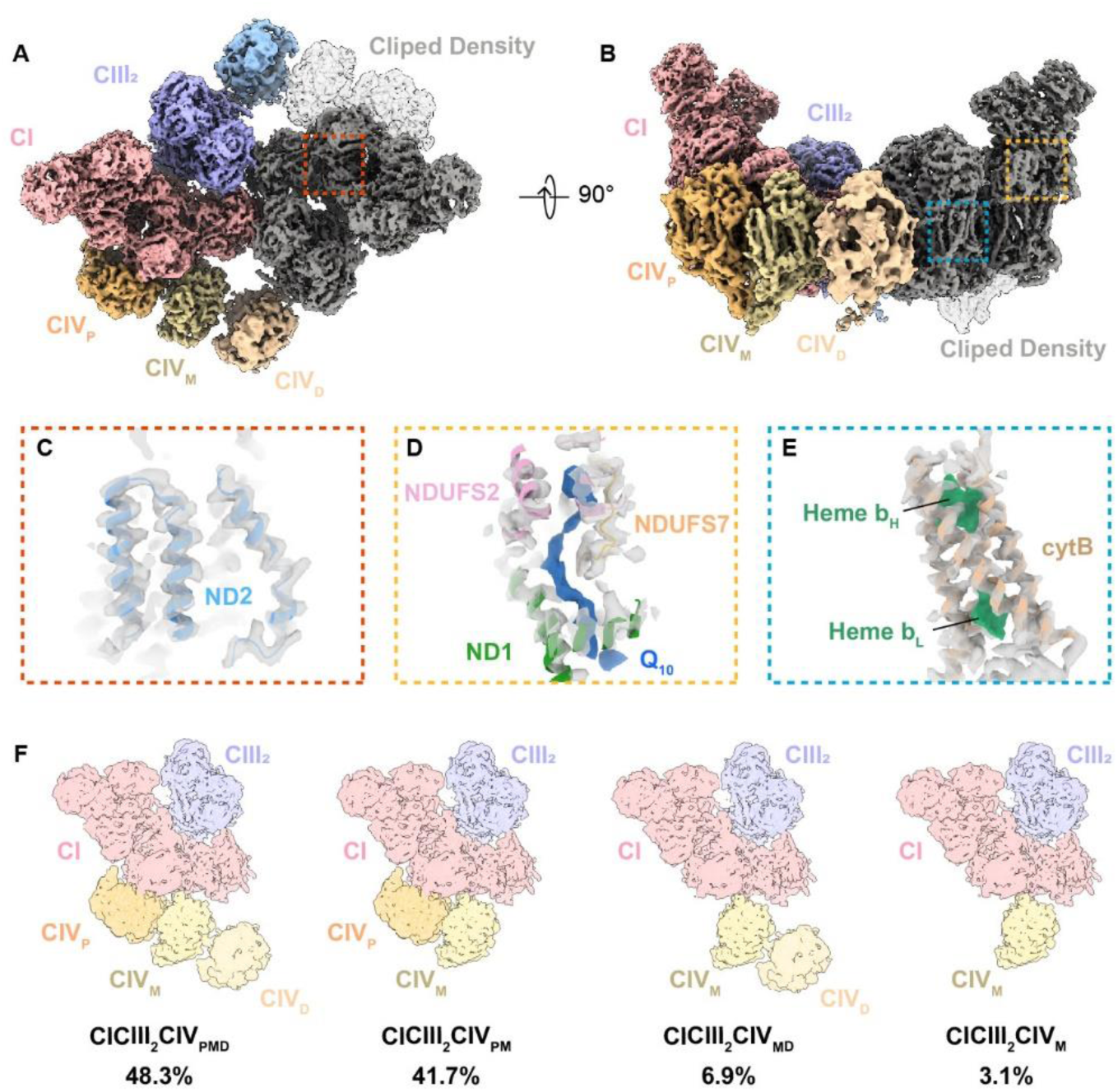
High-resolution in situ structures of respiratory chain complexes in *C. reinhardtii*. (**A, B**) Overall densities of the CI_2_CIII_4_CIV_6_ supercomplex viewed from the matrix side (**A**) and the membrane side (**B**). The CICIII_2_ template is colored dark gray, with differences from the purified structure (EMD-50202) shown in distinct colors. (**C–E**) Close-up views of representative features resolved in the respiratory chain complexes, with densities shown in gray. (**C**) Transmembrane side chains of the ND2 subunit, with the model colored in blue. (**D**) The active Q10-binding site and surrounding environment in CI, with the Q10 density shown in dark blue and the model of CI colored accordingly. (**E**) Heme groups in CIII, shown as green densities, together with surrounding cytB helices, with the model colored in brown. (**F**) Cartoon representations of four distinct compositional states of the CICIII₂CIV_n_ fundamental unit of respiratory chain complexes and their relative proportions.

### Rapid Capture of the Photoprotective State of PSII

Numerous essential biological processes difficultly resolved for SPA remain to be determined using in situ structural analysis, such as photoprotection in photosynthetic organisms: PSII is susceptible to oxidative damage under excess illumination, which is mitigated by rapid dissipation of excess energy as heat ^62^. The primary photoprotective process, energy-dependent quenching (qE), is activated and relaxed on a timescale of seconds to minutes ^63,64^, making structural determination particularly challenging. The formation and maintenance of the energy-dissipation state of PSII associated with photoprotective proteins depend on environmental stimuli, such as the light intensity. By integrating cryo-LEP-FIB and cryo-iSIA, we established the efficient strategy that enables structural determination of in situ macromolecules—from lamellae preparation to data processing—within 3 days. This capability allowed us to rapidly assess the photoprotective protein-associated state of PSII under different light conditions. In the absence of stimulation, the complex predominantly adopted a single, unassociated structural state. Upon stimulation, a distinct associated state became apparent and could be detected (Figure S17) in cryo-LEP-FIB–prepared lamellae. When stimulation was interrupted during vitrification, this association rapidly disappeared, indicating the highly transient nature of this functional state. Overall, this unified strategy enables rapid, high-resolution structural feedback, substantially reducing experimental iteration and accelerating the identification of optimal conditions for target proteins in defined physiological states.

## Discussion

It is now well established that cryo-FIB is a key technique for cryogenic thinning of biological specimens and in situ structural analysis. However, biological samples are considerably more sensitive than materials to ion implantation and secondary-electron damage ^26,65^, and therefore damage estimates derived from material systems usually cannot be directly applied to biological specimens. Moreover, cryo-FIB has a comparatively short history in biological sample thinning, and early studies were limited by resolution, likely underestimating the damage depth. In addition, low-energy milling of biological specimens was technically more challenging than conventional milling. These factors have therefore limited the widespread adoption of low-energy milling in cryogenic biological sample preparation. In this study, Cryo-LEP-FIB enabled the preparation of thinner, higher-quality cryo-lamellae and markedly improved isSPA localization, facilitating the successful determination of high-resolution structures of multiple in situ protein complexes. These results highlight the practical value of low-energy milling for biological specimen preparation and demonstrate its broader potential to advance high-resolution cryo-EM structural analysis. It should also be noted that all data in this study were collected using a K2 camera. Our unpublished data suggest that with a Falcon IV camera, the resolutions of in situ reconstructions could be further improved probably due to a better refinement with increased data quality.

High data quality is critical for in situ structural analysis, and continued improvement of thin-lamella preparation remains essential for broadening the analytical scope and improving the resolution of this approach. Given that the damage depth at 8 keV is 30 nm, the current lamella thickness of 60–80 nm already approaches the practical limit for preserving undamaged samples at this voltage. To further lower the molecular-weight limitation for structure determination by in situ single-image analysis, such as isSPA and 2DTM, it may be necessary to explore even lower cryo-FIB polishing voltages. However, practical implementation of cryo-FIB polishing below 5 keV in cryo-biological samples remains challenging due to physical limitations of the ion beam. Future progress will likely depend on improved ion-source designs that maintain sufficient beam current and focus at lower voltages, together with more efficient low-energy milling strategies for biological specimens. In addition, while 30 keV thinning can now be automated through software control, the subsequent low-energy polishing step still relies on manual operation. Developing a fully automated low-energy polishing workflow could greatly enhance thinning efficiency. Additionally, ultrathin lamellae below 60 nm are prone to wrinkling or fracturing during transfer, further limiting the feasibility of lower-energy milling. As a result, the transfer stability of ultrathin lamellae remains a major practical bottleneck.

Recently, plasma FIB (pFIB) instruments have emerged as promising tools for large-scale lamella production, as plasma ion species can achieve smaller probe sizes than Ga^+^ at high beam currents ^34,66^, thereby improving milling throughput. Previous studies have estimated that the highest-quality particles in pFIB-milled lamellae reside beyond 45 nm from the milling surfaces; however, the limited resolution of subtomogram averaging has prevented a more precise determination of the damage depth. In our study, the pFIB/SEM system available to us exhibited lower imaging resolution at low current and voltage than Ga^+^-source instruments, which prevented the preparation of sufficiently thin lamellae for a systematic evaluation of voltage-dependent damage. Given that pFIB is expected to have a beam-damage mechanism similar to that of Ga^+^-based FIB, low-energy polishing is also likely to be beneficial, and its higher milling rate may offer additional opportunities for optimizing lamella preparation.

Considering the limitations described above, obtaining large datasets from thinner, less-damaged lamellae, particularly those below 60 nm, remains challenging. Our data show that proteins around 400 kDa were efficiently detected within lamellae ≤120 nm thickness (Figure 3D), a thickness range of 60–120 nm is optimal for high-resolution in situ structural analysis of proteins of relatively low molecular weight. It is worth noting that target proteins often form higher-order assemblies with endogenous macromolecules in their native environment as cases shown above ^50,51^. Therefore, we recommend using thinner lamellae for efficient localization of low-molecular-weight proteins, followed by a second-round matching step in which using higher-order assemblies reconstructed from the target protein are used as template to search the complete dataset, including thicker lamellae, to maximize data utilization.

During isSPA processing, false-positive particle detections can introduce model bias, thereby limiting the resolution of in situ structures to around the frequency range used for particle detection. Novel densities within higher-order complexes can serve as valuable discriminative features in subsequent no-alignment 3D classification, helping distinguish true particles from false-positive detections and thereby reducing model bias. In PSII, for example, the peripheral antenna system surrounding the core provides useful structural features for classification. For proteins lacking sufficient novel densities relative to the template, model bias can be reduced by scoring at frequencies higher than those used for particle detection ^23^. In PSI, coarse classification followed by first-round refinement to 3.9 Å allowed particles to be sorted using a 4–6 Å frequency range to remove false-positive solutions.

In cryo-EM micrographs of cellular lamellae, densities from the crowded cellular environment overlap with those of the target protein and contribute background noise during particle localization. IsSPA is particularly sensitive to such background because it relies on a single 0° image for reconstruction. Thinner lamellae reduce overlapping densities and thereby improve particle localization. Accordingly, obtaining low-damage thin lamellae can markedly enhance the performance of isSPA. By contrast, cryo-ET reconstructs 3D volumes from tilt-series data, separating overlapping densities from the target protein along the z-axis. Particle picking in cryo-ET is performed through low-frequency 3D template matching, followed by iterative refinement to gradually improve resolution. This low-frequency strategy is largely insensitive to ion-beam damage, allowing even conventional 30-keV milled lamellae to support the localization of particles as small as nucleosomes (∼200 kDa) in very thin cell lamellae ^67–69^. In thin lamellae, however, the damaged layer occupies a larger fraction of the lamella, and the loss of SNR increasingly impairs the alignment of smaller proteins, thereby limiting high-resolution structure determination. Thus, it is possible that thin lamellae prepared by low-energy milling are particularly advantageous for high-resolution cryo-ET analysis of small proteins.

Although thinner lamellae are critical for in situ structural analysis, high-resolution structure determination is not the sole objective in the field. Equally important are the three-dimensional reconstruction of subcellular architecture and the spatial localization of proteins. These analyses require thicker lamellae (200–300 nm) to preserve native spatial context at the subcellular level. Tailoring technical approaches to specific research goals is essential—cryo-ET excels in visualizing the in situ organization and interactions of macromolecules within intact cellular environments, while cryo-iSIA offers enhanced efficiency and precision for resolving targeted complexes within crowded cellular milieus. Together, these complementary methods enable a coherent approach to in situ structural analysis.

## Data and materials availability

The associated electron density maps have been deposited in the Electron Microscopy Data Bank under the following accession codes: EMD-66522 (*C. reinhardtii* 80S ribosome), EMD-66521 (*S. cerevisiae* 80S ribosome), EMD-66520 (*E. coli* 70S ribosome), EMD-66518 (PSII, dimeric core template), EMD-66529 (CP26-S region of PSII), EMD-66528 (CP29-M-L region of PSII), EMD-66527 (PSII, monomeric core template), EMD-66526 (*C. reinhardtii* PSI), EMD-66525 (*C. reinhardtii* PSI-LHCII), EMD-66524 (LHCII region of PSI-LHCII), EMD-66523 (*C. reinhardtii* Rubisco), EMD-66519 (*C. reinhardtii* chloroplast ribosome), EMD-69200 (*C. reinhardtii* CI_2_CIII_4_CIV_6_ supercomplex), EMD-69057 (*C. reinhardtii* CICIII_2_CIV_3_ supercomplex), EMD-69059 (*C. reinhardtii* CI), EMD-69062 (*C. reinhardtii* CIII_2_), EMD-69061 (*C. reinhardtii* CIV_P_), EMD-69058 (*C. reinhardtii* CIV_M_),which are publicly available as of the date of publication.

## Acknowledgments

Cryo-EM data collection was carried out at the Center for Biological Imaging (CBI), Core Facilities for Protein Science at the Institute of Biophysics (IBP), Chinese Academy of Sciences (CAS). We thank X. J. Li, L. L. Qin, and other staff members at the CBI (IBP, CAS) for technical support in data collection. We thank L. F. Kong for cryo-EM data storage and backup. The project was funded by the National Key R&D Program of China (2024YFA1307402, 2024YFA1307303 and 2021YFA1301501), the National Natural Science Foundation of China (32150010, 32325027, 32401017 and 32241027), the Key Research Program of Frontier Sciences at the Chinese Academy of Sciences (ZDBS-LY-SM003) and Basic Research Program Based on Major Scientific Infrastructures, CAS (JZHKYPT-2021-05). Key Laboratory of Biomacromolecules, Chinese Academy of Sciences (ZGD-2023-05). CAS Project for Young Scientists in Basic Research (YSBR-106 and YSBR-015).

## Author information

These authors contributed equally: Chunling Wu, Qi Yang.

## Authors and Affiliations

**State Key Laboratory of Biomacromolecules, Institute of Biophysics, Chinese Academy of Sciences, Beijing 100101, China**

Chunling Wu, Qi Yang, Xiaodong Su, Mei Li, Xinzheng Zhang

**University of Chinese Academy of Sciences, Beijing, China**

Qi Yang

## Contributions

X.Z. supervised the project; C.W. and Q.Y. performed the experiments, including preparing the *S. cerevisiae* and *E. coli* cells, developing cryo-LEP-FIB methods, milling the cells to cellular lamellae, collecting the cryo-EM data, performing the data processing, and conducting the data analysis; X.S. and M.L. prepared the *C. reinhardtii* cells. All authors contributed to the writing of the manuscript.

## Corresponding authors

Correspondence to Xinzheng Zhang.

## Ethics declarations

### Competing interests

The authors declare no competing interests.

## Methods

### Cell culture and cryo-sample preparation

Cells of the *C. reinhardtii* CC124 strain were grown in TAP medium at 23 °C with shaking at 130 rpm under continuous illumination. For cryo-EM specimen preparation, all cells were diluted to 0.01 mg/mL chlorophyll in fresh TAP medium.

*S. cerevisiae* strains BY4741 were inoculated in YPD medium grown overnight at 29 °C to an OD600 of 0.6–0.8. The cells were collected and resuspended in the same medium, and the concentration of the suspension was adjusted to an OD600 of 2.

*E. coli* BL21 (DE3) cells carrying the kanamycin resistance marker were first grown overnight in LB medium supplemented with kanamycin at 37 °C with shaking at 220 rpm. The overnight culture was then diluted 1:100 into fresh LB medium and incubated under the same conditions until an optical density at OD600 of 0.6–0.8 was reached. Cells were harvested by centrifugation at 2000 ×g for 3 min, resuspended to achieve a 10-fold concentration, and frozen for subsequent use.

For cryo-EM specimen preparation, 3 µL of cells were applied to glow-discharged grids: quantifoil R1.2/1.3 Cu 200 mesh grids were placed horizontally for 30 s to allow even cell distribution before plunge-freezing in a 1:1 mixture of liquid ethane and propane using a Leica EM GP (Leica Microsystems) with 4–6 s back-side blotting at 16 °C and 70% chamber humidity.

### The workflow of cryo-LEP-FIB

Cryo-lamellae were prepared by cryo-FIB milling using a dual-beam cryo-FIB-SEM Aquilos 2 system (Thermo Fisher Scientific). Before milling, the grids were sputter-coated with metallic platinum for 15s using the beam current of 30mA then coated with organo-platinum for 60s through the GIS and coated metallic platinum for 15s again with the same settings. Because GIS designs and deposition efficiencies vary across different cryo-FIB systems, the organo-platinum deposition time was empirically adjusted. For cryo-LEP-FIB, extending the GIS deposition to approximately twice that used for 30 keV Ga⁺ milling performed on the same system provided robust surface protection and improved lamella stability during low-energy polishing. Rough milling at 30 keV was performed either manual or automated milling. For manual rough milling, stress grooves were milled ∼3µm away from the left and right side of target position using 30 keV 1.0 nA Ga^+^ beam. The lamella was then rough-thinned to ∼300 nm by sequentially reducing the Ga⁺ beam current from 0.5 nA to 30 pA (Figure 1B). Milling was carried out in Si mode with a milling angle of 10°, and the Z-axis milling depth was set to ∼0.8 µm. For automated rough milling, lamellae were prepared using Thermo Fisher’s AutoTEM Cryo 2.4 software, progressively reducing the Ga⁺ ion beam current from 1.0 nA to 50 pA at 30 keV, achieving an approximate thickness of 300 nm. After high energy rough milling, ion voltage was changed to 8 keV for low energy polishing, a 75pA was used to polish the lamellae to less 150nm. To achieve a more uniform lamella thickness, an increased over-tilt angle was applied relative to conventional 30 keV polishing (typically ±0.2°). Specifically, the over-tilt angle was set to 10 ±0.6∼0.8°, depending on the lamella length. Specifically, an angle of ±0.6°was used for lamellae shorter than 10 µm, ±0.7°for those between 10 and 20 µm, and ±0.8°for lamellae longer than 20 µm. The Z-axis value of Si mode was set to 4 µm in this step to avoid curtain and to obtain uniform thickness. Comparable milling conditions were readily transferable to lamella preparation of all three cell types.

### Cryo-EM data collection and processing Data collection of iSPA

Grids containing milled lamellae were loaded into a 300 kV FEI Titan Krios G3 microscope (Thermo Fisher Scientific, USA) equipped with a Gatan GIF K2 direct electron detector (Gatan Inc., Warrendale, PA). Data were collected in super-resolution mode at a nominal magnification of 105,000×, with a binned pixel size of 1.32 Å.

Images were acquired using beam-image shift data collection methods ^70^. An offset start angle, corresponding to the FIB milling angle, was applied to ensure that the lamellae were perpendicular to the electron beam ^23,60^. Each movie stack consisted of 50 frames, with a total dose of 50 e⁻/Å², a dose rate of 15.6 e⁻/Å²/s, and a defocus range of -0.8 to -1.2 µm.

### Structure determination of cytoplasmic ribosome in *C. reinhardtii*

From *C. reinhardtii* lamellae, a total of 1,878 micrographs were acquired in half a day. Motion correction and dose weighting were applied using MotionCor2 ^71^, and CTF estimation was carried out with CTFFIND4 ^72^ implemented in RELION ^73^. Of these, 408 micrographs were selected for particle detection using the LSU of the cytoplasmic ribosome of *S. cerevisiae* (EMD-38656) as a template in isSPA, details described as previous studies ^60^, resulting a 4.1-Å resolution map of cytoplasmic ribosome of *C. reinhardtii*. The LSU of *C. reinhardtii* was then extracted as a new template to identify as many 80S ribosomes as possible, with template projections generated at 3°angular increments. Particle detection was performed using an overlap parameter of 3 and a frequency range between 1/400 Å⁻¹ and 1/6 Å⁻¹, resulting in 208,508 particles that passed a score threshold of 6.5. After alignment-free 3D classification and refinement in RELION, 39,711 particles were selected and resulted a resolution of 3.6 Å of LSU and 4.0 Å of SSU. To further test the detection of smaller proteins, the body of SSU was extracted used as a template for particle detection with the same parameters as above. A total of 10,278 high-quality particles were refined locally, yielding an initial reconstruction at 4.2 Å resolution. Final CTF refinement in RELION followed by local refinement in CryoSPARC ^74^ improved the LSU map to 3.7 Å resolution (Figure S4 and Table S1).

### Structure determination of PSII in *C. reinhardtii*

A total of 892 micrographs of thylakoid membranes were selected for particle picking using the C_2_S_2_M_2_ region of PSII from *C. reinhardtii* (EMD-9956) ^52^ as the template in isSPA. Applying the same parameters and processing workflow described above yielded a C_2_S_2_M_2_L_2_-type PSII supercomplex at 3.9 Å resolution. The C_2_ region was then extracted as a new template to assess the data quality of lamellae prepared by cryo-LEP-FIB. Using a score threshold of 6.5, 332,356 particles were identified for subsequent alignment-free 3D classification. Classes containing LHCIIs were selected for another round of alignment-free 3D classification with 2 classes in CryoSPARC ^74^. A total of 57,382 particles were selected and used for local refinement, resulting in a 3.9 Å reconstruction of the C_2_S_2_M_2_L_2_-type PSII supercomplex. A round of CTF refinement and C2 symmetry expansion was performed, yielding an overall resolution of 3.4 Å with a C_2_S_2_M_2_L_2_ mask. Local refinements of the CP26 and S-LHCII regions yielded a 3.3 Å map. Another round of 3D classification was then performed in CryoSPARC using an M- and L-LHCII mask. From this, 73,219 particles (after C2 symmetry expansion) with more continuous densities were selected for local refinement, yielding a 3.4 Å resolution map of the M- and L-LHCII region (Figure S8).

Using the 400-kDa monomeric core of PSII as the template, 576 micrographs with a thickness below 140 nm were selected for particle detection using identical parameters. Particles with a matching score above 6.5 were extracted and subjected to two rounds of 3D classification, sequentially performed in RELION and CryoSPARC. This process yielded 12,837 C_2_S_2_M_2_L_2_-type PSII particles, resulting in a 3.7 Å reconstruction. Subsequent CTF refinement and C2 symmetry expansion improved the overall resolution to 3.3 Å (Figure S9).

### Structure determination of PSI in *C. reinhardtii*

The same 892 micrographs of thylakoid membranes were used for the in situ structural determination of PSI. Using the PSI complex (EMD-9678) ^54^ as the template, 313,106 particles were identified using the same particle detection parameters as described above, with a score threshold of 6.5. Particles were extracted with a box size of 360 pixels. 3D classification in CryoSPARC was then performed, and the class showing additional densities corresponding to Lhca2 and Lhca9 was selected. A subsequent round of local refinement yielded a 3.9 Å reconstruction of the PSI supercomplex. To further reduce template bias, the isSPA sorting algorithm ^22^ within the resolution range of 4-6 Å was applied, and particles with scores below 0.012 were discarded, resulting in 56,000 high-confidence particles. After one round of CTF refinement, the overall resolution improved to 3.3 Å. To resolve conformational heterogeneity, another round of 3D classification was conducted in CryoSPARC, revealing two major classes: the PSI–LHCII supercomplex (25.1%) and the PSI complex without LHCII (74.9%). Local refinements yielded reconstructions at 3.8 Å and 3.3 Å resolution, respectively. The local resolution of the LHCII region in the PSI–LHCII supercomplex reached 4.7 Å (Figures. S11 and S12).

### Structure determination of Rubisco in *C. reinhardtii*

105 micrographs containing chloroplast pyrenoids were selected for the in situ structural determination of Rubisco. Particle detection was performed using the cryo-EM map of Rubisco (EMD-22308) ^58^ as the template in isSPA, with template projections generated at 3° angular intervals under C1 symmetry. The matching frequency parameters were the same as described above, and particles with scores above 6.2 were selected, yielding 180,782 candidates. To remove redundancy, particles within 6 Å of each other were considered duplicates and excluded. After applying D4 symmetry expansion, 627,192 particles were obtained. A round of 3D classification was then carried out in CryoSPARC, and the high-confidence class was selected based on the presence of extra densities corresponding to EPYC1 and the complete N-terminal domain of the LSU. Following local refinement and CTF parameter optimization, the final reconstruction reached an overall resolution of 3.2 Å from a total of 188,820 particles after symmetry expansion (Figure S13).

### Structure determination of Chloroplast ribosome in *C. reinhardtii*

Using a 6.2-Å map of the chloroplast ribosomal LSU (a gift from the Li lab.) as the template, particle detection was performed in isSPA with frequencies ranging from 1/400 Å⁻¹ to 1/8 Å⁻¹, yielding 363,033 particles. Two rounds of alignment-free 3D classification without a mask, followed by one round with an SSU local mask, identified 29,867 particles for further refinement. To further remove biased particles, the isSPA sorting algorithm was applied within the resolution range of 4-6 Å. Particles with scores below 0.027 were discarded, leaving 11,588 high-confidence particles. After CTF refinement, the chloroplast ribosome was reconstructed at an overall LSU resolution of 4.0 Å (Figure S14).

### Structure determination of mitochondrial respiratory chain complexes in *C. reinhardtii*

A total of 910 micrographs containing mitochondria were selected for particle picking using a purified CICIII₂ complex from *C. reinhardtii* (EMD-50202) ^11^ as the template in isSPA, with template projections generated at 3° angular increments. Particle detection was performed over a frequency range of 1/400 to 1/6 Å⁻¹, yielding 380,719 particles that passed a score threshold of 6.5. Following alignment-free 3D classification in RELION, 54,073 particles were retained and recentered using block-based reconstruction (BBR) ^75^. Subsequent classification in CryoSPARC identified a CI₂CIII₄CIV₆ respiratory chain supercomplex. Redundant particles within 80 Å were then removed. After C2 symmetry expansion and another round of alignment-free 3D classification focus on the CICIII₂CIV₃ asymmetric unit, 17,535 particles were selected. CTF refinement yielded an overall resolution of 4.08 Å for the CI₂CIII₄CIV₆ supercomplex. Local refinement of the asymmetric unit, resolved the CICIII₂CIV₃ supercomplex at 3.85 Å resolution. Further local refinements of the CI, CIII₂, and CIV_M_ regions yielded maps at resolutions of 3.7 Å, 3.85 Å, and 4.1 Å, respectively. Additional classification using a CIV_P_ mask revealed that 15,786 particles (∼90%) contained CIV_P_, whereas a minority class of 1,749 particles (∼10%) lacked CIV_P_. Further focused classification on CIV_D_ complex based on CIV_P_-occupied particles separated two classes corresponding to particles with and without CIV_D_, comprising 8,475 and 7,311 (Figures. 5F and S15J) particles, respectively. Subclassification of the CIV_P_-unoccupied particles on CIV_D_ complex identified two distinct states: one containing both CIV_M_ and CIV_D_ (1,218 particles), and another containing CIV_M_ only (531 particles, Figures. 5F and S15I).

### Structure determination of cytoplasmic ribosome in *S. cerevisiae* and *E. coli*

347 micrographs of *S. cerevisiae* and 229 micrographs of *E. coli* were selected. Particle detection was performed in isSPA using the cryo-EM ribosome maps EMD-38656 ^60^ (*S. cerevisiae*, SSU body) and EMD-22586 ^76^ (*E. coli*, SSU body) as templates (Figures. S5 and S6), with projections generated at 3°intervals under C1 symmetry. Candidates were selected with scores above 6.2, yielding 90,290 (*S. cerevisiae*) and 647,004 (*E. coli*) particles. After two rounds of non-alignment 3D classification in RELION (the second iteration with an LSU mask), high-confidence classes were identified by the presence of LSU densities. Subsequent local refinement (LSU mask) and CTF parameter optimization yielded reconstructions of the LSU at 3.4 Å (*S. cerevisiae*, 20,604 particles, Figure S5) and 3.5 Å (*E. coli*, 8,691 particles, Figure S6).

### Calculation of P-R curves

The calculation of the P-R curve followed our previous study ^22,23^ and is briefly described below. Precision was defined as the ratio of true particles to all particles detected by isSPA, and recall as the ratio of true particles to the ground truth. For ribosomes in the three cell types, the ground truth was defined as the particles identified by applying the LSU from the 3.6-Å cryo-EM map of *C. reinhardtii* (described above), EMD-38656 (*S. cerevisiae*), and EMD-22586 (*E. coli*) as templates for particle detection and alignment-free 3D classification. Blocks of different molecular weights were extracted from the corresponding LSU densities. For PSII in C. reinhardtii, the ground truth was defined as the particles identified using the C_2_S_2_M_2_ region of cryo-EM map EMD-9956 as the template for particle detection and alignment-free 3D classification, while blocks of different molecular weights were extracted from the C2S2M2 region of the in situ PSII reconstruction. All datasets in this study were collected using the same cryo-electron microscope as in our previous studies. Thus, the relative lamella thickness was comparable and estimated using the Beer-Lambert law ^23,26,77,78^. For P-R curve calculations, 10-20 micrographs within the same thickness range were selected every ∼20 nm.

### Detection efficiency analysis by isSPA across lamella depth layers

Particle detection efficiency across different lamella depth layers was analyzed using a cryo-lamella dataset of concentrated virus samples from a previous study ^26^. Briefly, concentrated parvovirus particles were plunge-frozen on EM grids and milled using an Aquilos-2 cryo-FIB/SEM microscope at either 8 kV or 30 kV. For data collection, a high-dose single-particle image was first acquired with the lamella oriented horizontally, followed by acquisition of a tilt series for cryo-ET reconstruction. Particle coordinates in the X-Y plane were obtained from the single-particle image via isSPA-based detection. Corresponding Z coordinates (depth from the lamella surface) were determined from the tomographic reconstruction by aligning the X-Y coordinates of particles between the single-particle image and the tomogram. Particles were then assigned to depth layers of 10–20 nm, 20–30 nm, 30–40 nm, 40–50 nm, and 50–60 nm based on their Z coordinates. Particle picking for isSPA was performed using two templates: (i) the full virus density, with small regions excluded to minimize model bias, which was used to define ground-truth particles; and (ii) a clipped ∼400 kDa block derived from the virus, which was used for detection efficiency analysis. Recall was defined as the fraction of ground-truth particles that were correctly detected using the ∼400 kDa block template (Figure S8).

## Notes

### Competing Interest Statement

The authors have declared no competing interest.

